# Uncovering Co-regulatory Modules and Gene Regulatory Networks in the Heart through Machine Learning-based Analysis of Large-scale Epigenomic Data

**DOI:** 10.1101/2023.04.28.538783

**Authors:** Naima Vahab, Tarun Bonu, Levin Kuhlmann, Mirana Ramialison, Sonika Tyagi

**Author notes:** Corresponding author: Sonika Tyagi.

## Abstract

The availability of large scale epigenomic data from different cell types and conditions has provided valuable information to evaluate and learn features that predict co-binding of transcription factors (TF). However, previous attempts to develop models for predicting motif cooccurrence were not scalable for global analysis of any combination of motifs or cross-species predictions. Further, mapping co-regulatory modules (CRM) to their gene regulatory networks (GRN) is crucial in understanding the underlying function. Currently, there is no comprehensive pipeline to locate CRM and GRN on a large scale with speed and accuracy. In this study, we analyzed and evaluated different TF binding characteristics that would facilitate co-binding with biological significance to identify all possible clusters of co-binding TFs. We curated the UniBind database, which contains ChIP-Seq data from over 1983 samples and 232 TFs, and implemented two machine learning models to predict CRMs and potential regulatory networks they operate on. We narrowed our focus to study heart related regulatory motifs. Our findings highlight the importance of the NKX family of transcription factors in cardiac development and provide potential targets for further investigation in cardiac disease.

## 1 Introduction

Epigenomics provides insight into gene expression regulation. Among the many assays used in epigenomic studies, DNase-seq [1], ATAC-seq [2], and chromatin immunoprecipitation (ChIP) assays [3] are the most commonly used techniques. Of these listed methods, ChIP technology provides higher resolution and less noisy data which capture DNA fragments containing the position of transcription factors (TFs) binding or histone marks. Publicly accessible databases of ChIP-seq data include ENCODE [4], Unibind [5], and ChIP-Atlas [6].

Computational tools are used to identify TF binding sites (TFBSs) from ChIP-seq data, which are important for understanding gene regulation. Aligned profiles of TFBSs are called *cis*regulatory motifs. Popular motif finding tools include ChIP-exo [7], MEME [8], and HOMER [9]. These methods focus on searching for an individual binding site at a time and do not account for cooperation between the binding sites. Additionally, Deepbind [10], FactorNet [11], and BPNet [12] are machine learning approaches based on Deep Neural Network (DNN) to discover patterns of individual motif occurrences even without any labeled sequences. Although such neural network approaches require a considerable amount of processed data and enormous training, once the model is trained, it performs motif finding tasks with high specificity [13]. Deepbind models were trained on 12 terabases of sequence data while BPNet was trained on 147,974 genomic regions. FactorNet trains a DNN on data from one or more reference cell types for which TFs of interest have been profiled, and this model can then predict motifs in other cell types. Another approach, TF-MoDISco [14] uses partially redundant information in DNN to predict a non-redundant set of motifs learned by the convolutional filters.

Since multiple TFs are involved in regulating expression of a gene, identifying clusters of TFBS are critical in elucidating gene regulation. These clusters are referred to as *cis*-regulatory modules (CRMs). Each module is typically 300 bp or more in length and contains on the order of ten or more binding sites for at least four unique TFs [15][16]. This is the definition of CRM we will use in the rest of the article.

A few algorithms to identify CRMs from ChIP-seq data have been published, such as MCOT [17], GEM [18][19], TACO [20], and TF-Cluster [21]. The Motifs Co-Occurrence Tool (MCOT) uses a permutation approach to find one or more suitable partner motifs for the first motif occurrence. MCOT approach relies on over-representation from a single ChIP-seq experiment and does not consider cooperative binding of multiple proteins. Furthermore, MCOT is a species-specific method and works only for three species: *Homo sapiens, Mus musculus*, and *Arabidopsis thaliana*. Next, TF-Cluster generates a cluster of TF motifs based on shared co-expressed genes between multiple TFs. However, its dependency on available gene expression data limits is application in a genome-wide search. The GEM and TACO approaches were trained on known ChIP-seq data from 63 TFs and 29 TFs, respectively, to predict new motif co-occurrences. Hence, they are limited in their prediction accuracy and predict clusters of only two motifs. More recent and relevant algorithms to identify CRMs are BICORN [22], regCNN [23], dePCRM2 [24], and BPNet. BICORN uses a Bayesian-based approach along with TF-gene binding events and gene expression data as input to find a list of candidate CRMs. The regCNN algorithm extracts data from 5 different resources related to TF binding motifs, nucleosome free and variant sites, chromatin binding protein targets, histone modification information, and conservation score data, and feeds it into a CNN to predict whether a given chromosomal location contains CRM or not. Another tool, dePCRM2, extracts binding peaks from ChIP-seq and associated expresison, enhancer and variation informaiton from different dataset. It then constructs a graph with TFs as nodes. It adds an edge between two nodes if the corresponding TFs interact with each other based on the interaction information from known pathways. This algorithm uses the node perception program in *Cdlib* to cut the graph into smaller communities with 3 TFs in a CRM. In addition to these methodologies, the ChIP-Atlas database is comprised of peak calls from known ChIP-seq data and links up two ChIP-seq datasets for probable co-localized motifs. The platform provides the facility to visualize peak intensities of these co-occurring motifs. However, the data needs to be further analyzed to convert peaks into motif sequences along with their specific genomic coordinates.

All of the discussed methods either rely on known biological features to predict CRMs, and cannot provide predictions for TFs where corresponding feature datasets are ambiguous or unavailable. Further, these methods are designed to predict CRMs of short fixed lengths (up to 7). Importantly, none of the existing approaches connect the predicting of regulatory motifs to gene regulatory networks (GRN) they operate on.

GRNs describe interactions between genes and gene products, representing a collection of regulators that communicate with each other to govern gene expression levels that determine cell function [15][16]. GRNs can be used to derive novel biological hypotheses about molecular interactions and can be represented as graphs, with genes acting as nodes and inputs being proteins, such as transcription factors. Pathway models are important tools in systems biology to help understand complex biological systems made of GRNs [25] [26] [27].

In this study we have extended ChIP-seq data analysis for the prediction of CRMs. By studying TF co-binding and we developed a flexible, generic, species-agnostic method that can be used with or without prior feature generation to predict any combination of motifs. Our method allows mapping of the clusters back to GRNs in order to discover the TFs associativity and also report new TFs and genes nodes. We demonstrate application of our pipeline in cardiac case studies.

## 2 Methodology

### 2.1 Data Resources and Preprocessing

#### 2.1.1 Human genome annotations

We used human gene annotation in gene transfer format (GTF, version GRCh.38.p7) from the ENSEMBL database [28]. Chromosome names were renamed for consistency and random and unlocalized chromosome fragments were ignored. The dataset was then filtered to prepare a dataset containing 57955 genes, which is referred to as “gene data” in the article.

#### 2.1.2 TFBS recovered from ChIP-seq data

We used ChIP-seq captured bindings sites available via the Unibind database [5]. This database contains data from 1983 human ChIP-seq samples generated by over 11,000 ChIP-seq experiments from 9 different species. Although we only used human data for this study, the same workflow can be applied to data from other species. We processed the data to identify binding sites for a total of 321 motifs across “gene data”. For each motif, we combined multiple files of experimentally known binding sequences, removing duplicates and calling the resulting data “motif data”.

To study co-occurring motifs with biological significance, we focused on the promoter region, which is about 1500 base pairs (bp) upstream of the gene start position. Based on the literature [29], we relaxed the average distance between known pairs of motifs from 21 bp to 30 bp to account for any unknown variation (Figure S1). Pairwise combinations of the binding sites for the 321 motifs in the promoter region were generated. We further binned pairwise combinations for less than 1k bp, 1k bp to 5k bp and 5k to 10k bp to study the pattern of distance between motif pairs. We visualized the pairwise distances for less than 1500 bp and found that they had a skewed distribution with a mode of 9 bp (Figure S2), while distances greater than 1500 bp showed a uniform distribution(Figure S3).

As described above, we used ChIP-seq experiments to obtain positive data for motif pairs that met our constraints, while negative data was generated by considering motif pairs more than 1500 bp apart and not occurring in the promoter region of any gene. Both sub-datasets were combined and this resulted in a balanced data for co-occurring and non-co-occurring motif pairs.

### 2.2 ML Model Building for TF Co-occuring Prediction

The dataset was shuffled remove any ordering bias and combinations of co-occurring and non-cooccurring pairs was used to train two models: a CNN-based deep learning model and a Random Forest Classifier (RFC) model. The CNN model requires only TFBS sequences for prediction while the RFC model requires comprehensive data features for pairwise prediction. The CNN model simplifies initial input data, while the RFC model aids in feature selection and understanding of the most effective features for co-occurrence.

#### 2.2.1 Using Convolutional Neural Network (CNN)

CNNs excel in handling data with spatial or temporal relationships, represented as matrices. They employ convolution for feature extraction and pooling to curb dimensionality and prevent overfitting. Flattening layers convert features into vectors, acting as a bridge between convolution and dense layers. Additional layers like Dense and Dropout are also employed. In bioinformatics, CNNs have been valuable for DNA sequence analysis, especially in TFBS discovery. They efficiently search for short motifs within lengthy DNA sequences presented as matrices, eliminating the need for explicit feature extraction.

To provide the model with additional biological context, 30 base pairs (bp) of flanking sequences were extracted for each binding motif sequence. Due to a limited supply of labeled examples of longer clusters, a minimum cluster size of two was enforced. The binding sequences and their corresponding flanking sequences were aligned side-by-side with padding to accommodate different motif sequence lengths, creating a matrix format. Each pair of binding sequences was converted into an image format, and labeled examples were used for training. Hyperparameters were optimized on a representative 30% subset of the entire dataset. The final model was trained on the full dataset, which was split into 60:20:20 train:validation:test sets with 600K data points in the initial training set and 150K data points in the test set (Figure 1). Validation set was used to optimize the model hyperparameters.

**Figure 1:**
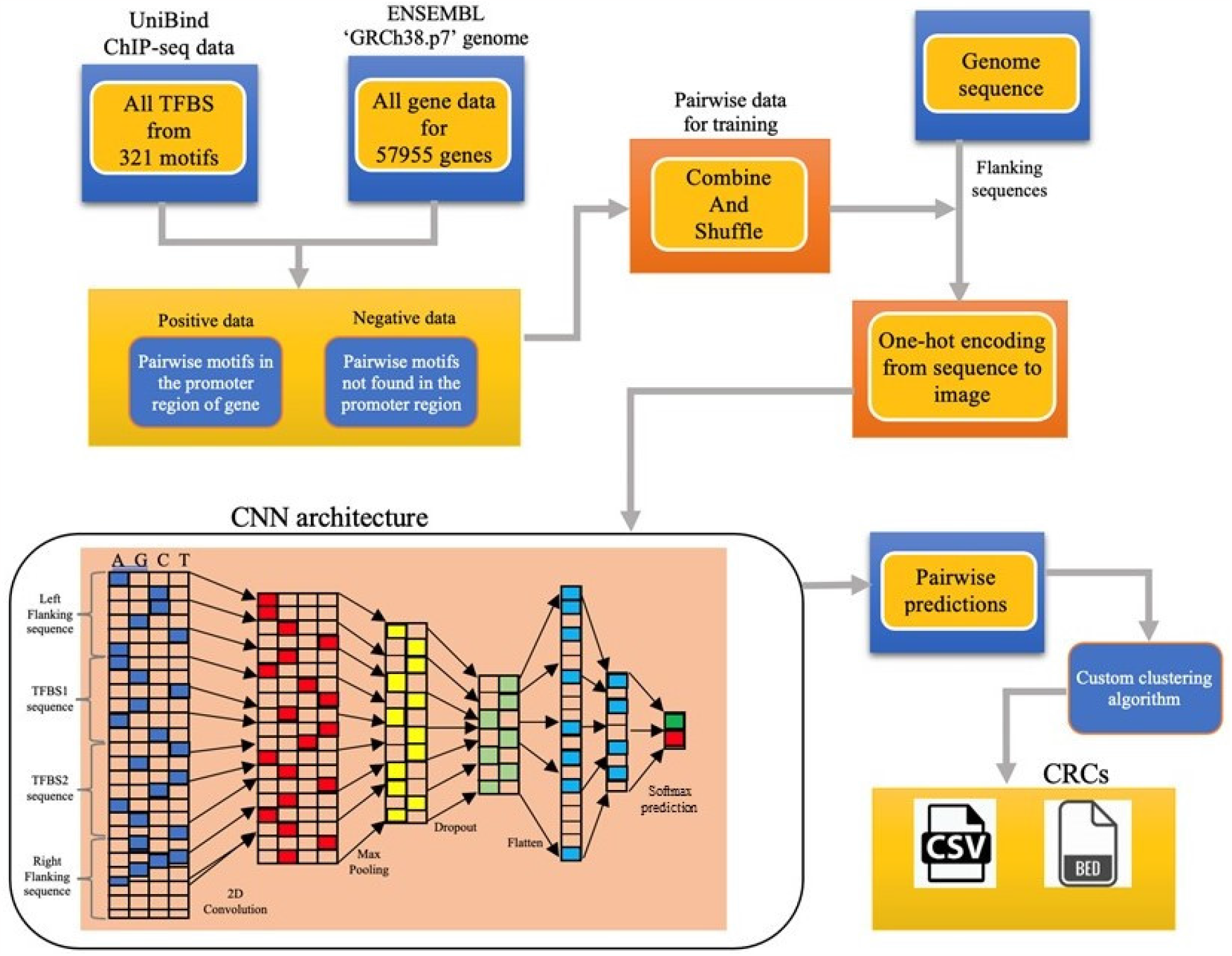
CNN Architecture and pipeline diagram for TF co-binding prediction

The final model consists of convolution layers with a number of filters ranging from 32 to 260, each with a kernel size of 4 * 4. These filters slide over the input vector to generate feature maps by applying convolution operations. The sub-sampling of the feature maps is performed using pooling layers. Here, we use a pooling layer that aggregates features by finding the maximum values in the filters. As the network progresses, the area of focus gradually reduces. Ultimately, the network can return the locations of the filtered sequences. The final model structure for the data is presented in Table 1, with a best validation accuracy of 0.92.

**Table 1:** Hyper parameters, range, and final Value used for the CNN model.

| Hyper parameter | Value Range | Final value |
| --- | --- | --- |
| Number of convolution layers | 3, 5, step=1 | 5 |
| Number of filters in convolution layers | 32, 260, step=32 | filters: 32, 256, 160, 192, 32 |
| Kernel size in convolution layers | (3, 5) | 4, 4, 4, 3, 4 |
| Activation function | ['relu', 'tanh'] | relu |
| The number of Dense layers | 1 to 5 | 2 |
| Number of Dense neurons | [32, 64, 128, 256, 512] | 512, 128 |
| Batch size | [64,200,600,1024,2000,3000] | 3000 |
| Drop out | [0.0,0.1,0.2,0.3,0.4,0.5] | 0.5 |
| Optimizer | ['adam', 'SGD', 'rmsprop'] | adam |
| Learning rate | [0.1, 0.01, 0.001,0.0001] | 0.0001 |

#### 2.2.2 Using Random Forest Classifier (RFC)

RFC is a machine learning algorithm that creates multiple decision trees and combines their predictions to make a final classification decision. This ensemble strategy helps RFC achieve accurate predictions and better generalization. RFC also calculates the importance of each feature, and the Gini feature importance method was used to study various features of DNA sequences that drive cooperative binding of TFs.

We base our approach on the assumption that DNA sequences contain short nucleotide patterns called “hotspots” that have biological significance. Some of these hotspots have TFBS and exhibit a nucleotide skew with polarity at the center of the consensus motif [30]. Nucleotide skew values can be calculated to quantify these hotspots, with the AT and GC skews being common ways to indicate over- or under-abundance of nucleotides in a sequence. Additionally, DNA shape features are the other set of features that can help predict the binding of certain TF families. Researchers have identified certain sequence and structural features that create a favorable microenvironment for binding [31] [32] [33]. As a result, these features can be studied in isolation or in combination with others to predict single TF binding motifs. Pre-computed protein-protein interaction (PPI) values are available for many TFs that can also be used as predicting features. Further, binding sites interact specifically with the DNA-binding domains of a TF, and hence, can be investigated for their role in motif identification.

In total we computed 43 features available for TF co-occuring prediction and by performing a recursive feature elimination with cross-validation, we found that 10 of these features are enough for optimal model training (Figure S4). Below we provide a detailed description of the features used in our model.

##### Nucleotide skews

The transcription factor binding site sequences were extracted for every pairwise combination to generate AT and GC skews [Equation 1] for *binding_site1* and *binding_site2* sequences. We also concatenated the sequences of both motifs to generate a combined skew.

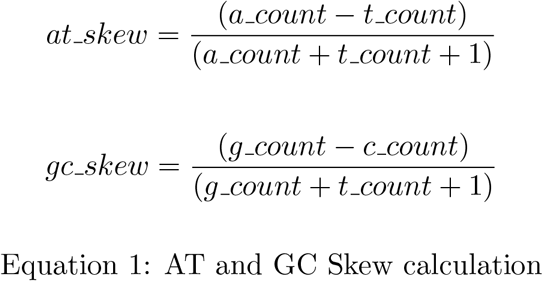

##### DNA shape scores

A shape score is assigned to each nucleotide in a DNA sequence, and these scores can provide insights into the readout modes of DNA sequences. We used 13 DNA shape features in our workflow as previously calculated by Chiu *et al* [34]. The sum of the respective scores for a 30 bp window centered at the motif sequence were calculated as the structural profile of the binding sites. The extraction of DNA shape scores was performed on the Monash eResearch High-Performance Computing (HPC) system using index-based Unix commands to retrieve precise scores at the motif’s location.

##### Protein-protein interaction score (ppi score)

The protein-protein interactions among TFs were obtained from the STRING database [26]. These interaction scores were combined and added to the dataset as a new feature called *ppi score*.

##### DNA binding domain (dbd)

The DNA binding domains (DBDs) of the TF binding motifs were categorized based on pre-determined domains obtained from the Protein Data Bank [35]. These domains were identified for every motif that was under consideration. This categorical feature was one-hot-encoded [36] before using in the model.

##### Frequency ratio (fr)

The frequency ratio measures the importance of motif pairs out of all possible combinatorial pairs. The score is calculated using the formula shown below [Equation 2].

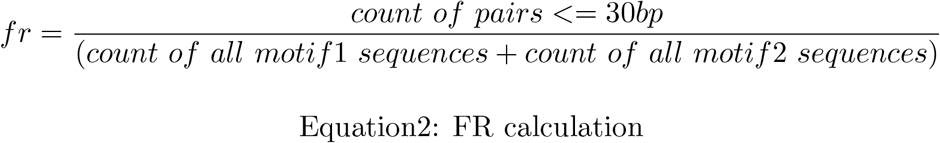

##### General motif features: motif1, motif2, chromosome, ng and dng

In addition to the biophysical features, we also recorded general representative features for the motifs. Since proteins select binding sites in a site-specific manner, a favorable microenvironment is required for TF binding to regulate their target genes [37]. Therefore, we extracted he distance between the binding sites (*dbbs*) and the distance to the nearest gene from the anchor motif (*dng*). Others were generic motif features such as the nearest gene (ng) name and its chromosome name for each motif pair (*motif1* and *motif2, chromosome*, and *ng*).

##### Target Variable: binding

The target variable is binary and is either ‘co-occurring’ or ‘non-co-occurring’. The positive dataset is labeled as co-occurring while the negative dataset is labeled as non-co-occurring. The features extracted from the dataset will be used to train the classification model.

Motif pairs without a known protein interaction score were kept separately as the “discovery set” to test for new co-occurring pairs.

### 2.3 Clustering and generation of GRNs

As the next step, a clustering algorithm was implemented to build biologically significant CRMs from predicted co-occurring motif pairs. The algorithm employed a ‘seed and extend’ approach where each motif pair with *motif1* or *motif2* is recursively extended by considering either motif1 or motif2 as seed motifs. The extended motifs were selected based on predicted co-occurrence and formed a cluster. This process generated a set of CRMs located in the promoter region of a gene. Sub-clusters were removed to extract a final CRM set.

#### Algorithm 1

Clustering algorithm

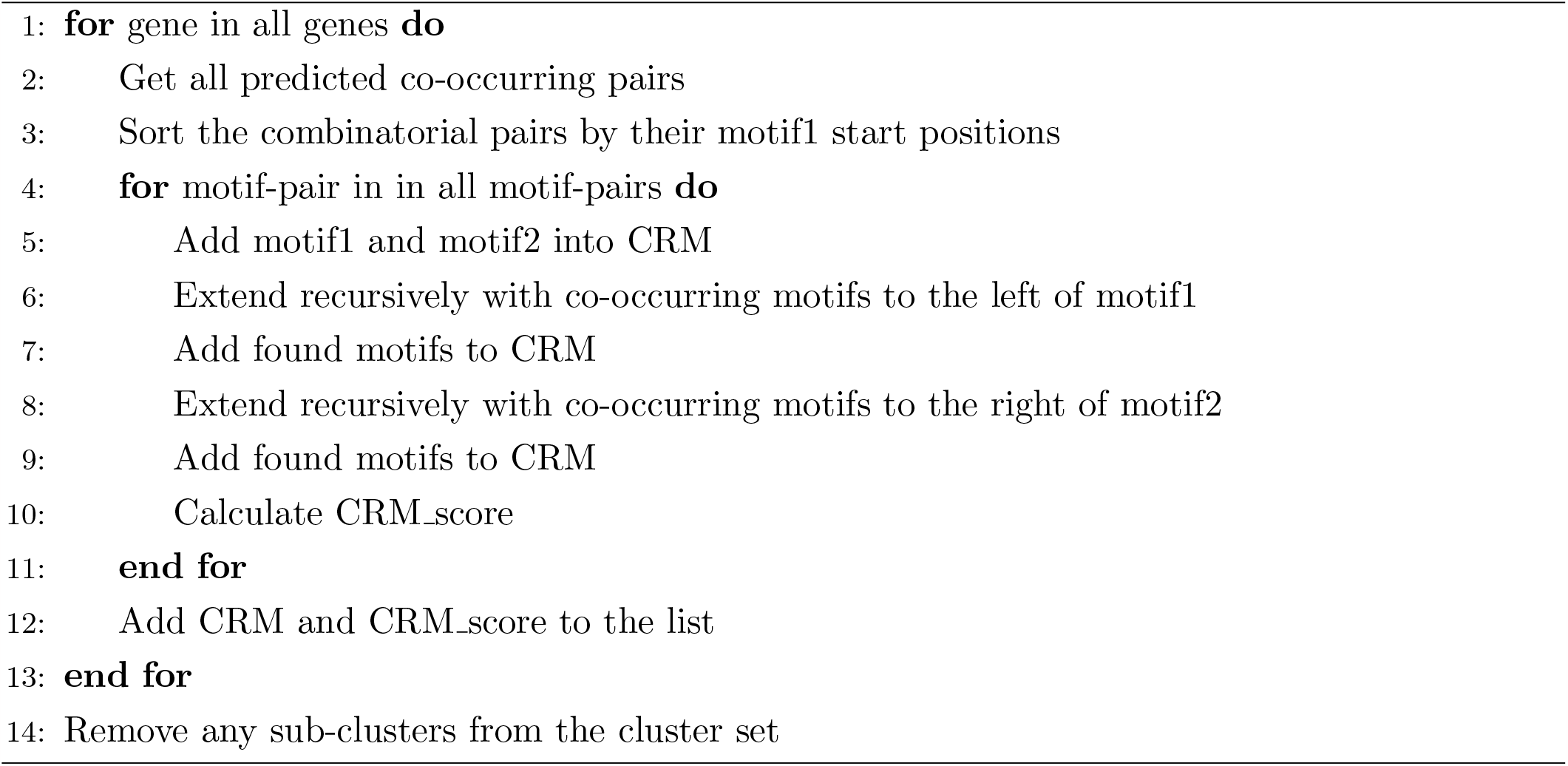

### 2.4 Finding gene regulatory networks (GRN)

After clustering, each gene will have multiple clusters corresponding to each TF binding. We only report clusters that have at least four different TFs and provide information such as the number of binding sites (some TFs can have multiple binding sites), binding site span, unique motifs, and their CRM score. We assume that genes sharing common motifs within their clusters form a regulatory network, which we visualize using a graph representation. By generating multiple graphs for the same genes, we can identify new motif nodes. We extract the list of genes from these graphs and use a pathway enrichment tool to map them to known biological pathways.

## 3 Results

### 3.1 TF Feature Importance

We used all 43 sequence-based, structure-based, and interaction-based features (Figure 2) that were scaled with a MinMax scaler[38], and RFC was trained on these features with 100 decision trees, determined through hyperparameter tuning. To understand the rules of cooperative binding, we also looked at individual features and their contribution to classify a motif cluster.

**Figure 2:**
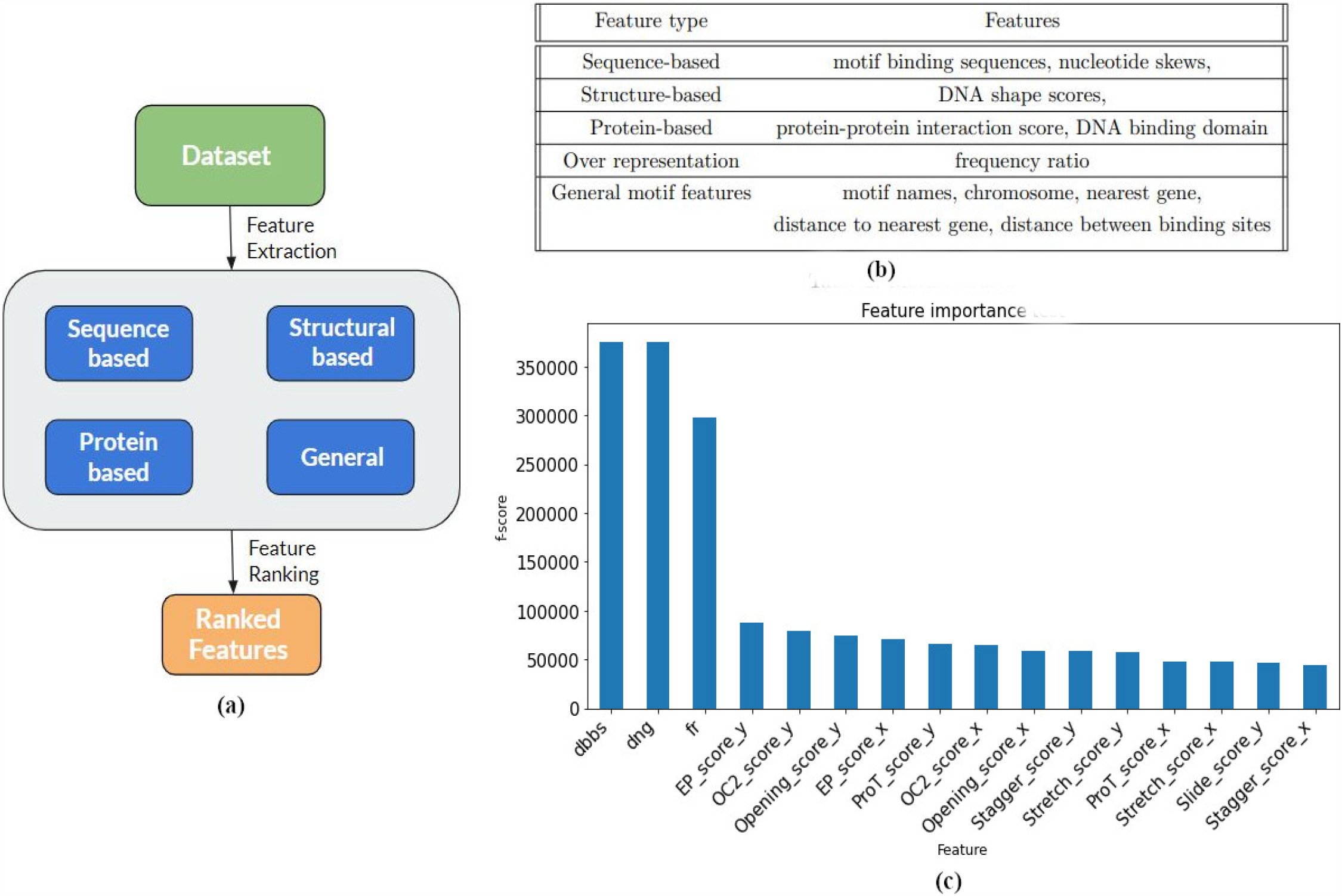
Feature Extraction Pipeline (a) The chromosomal dataset is preprocessed and different categories of features such as sequence-based, structural-based, protein-based and general features are extracted to input into a RF Classifier. (b) Features under each category is listed in this table (c) Once RF model is ready the features are ranked based on their importance score in order to model the predictions (see supplementary section *8*.*2* for more details)

**Figure 3:**
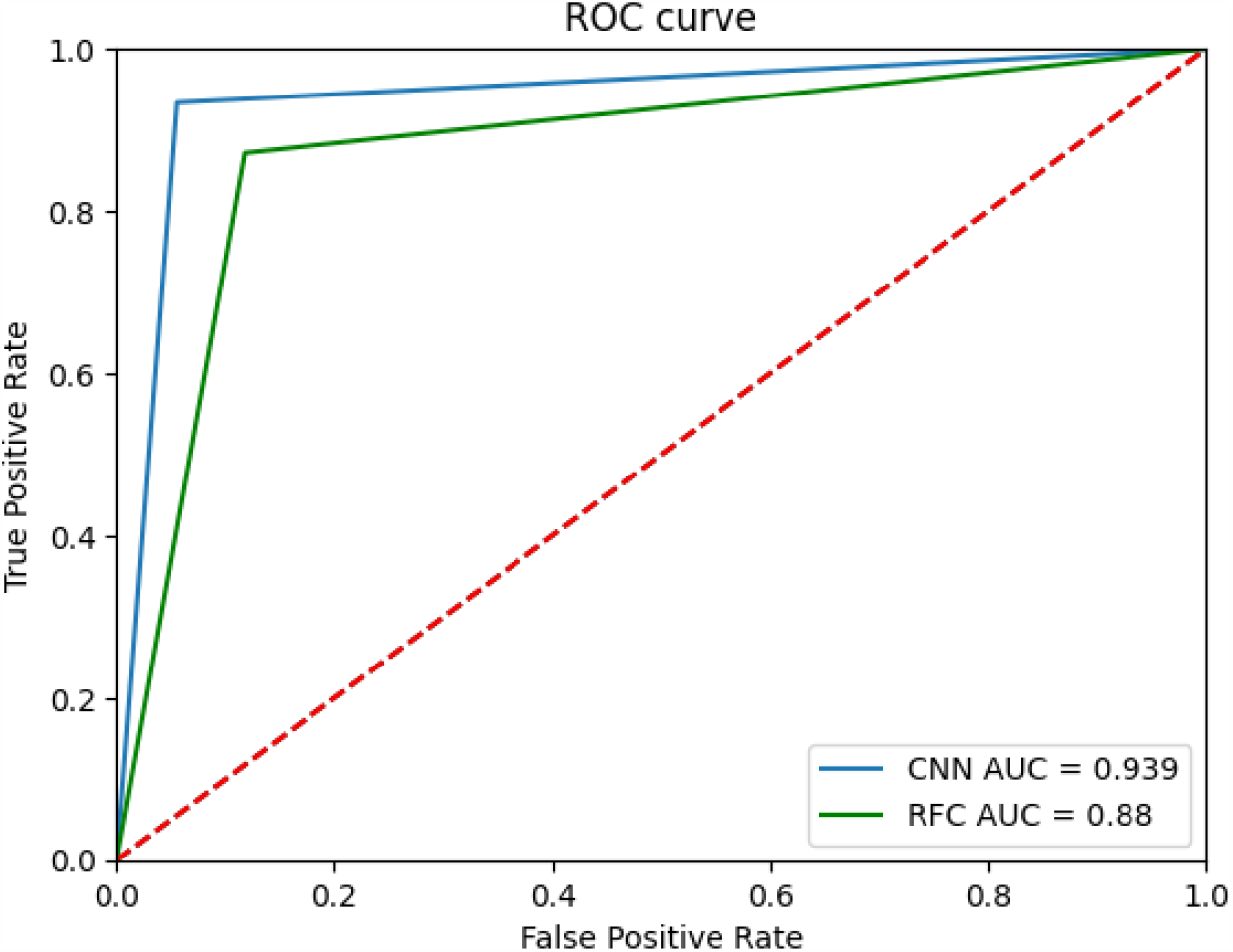
A Precision-Recall Receiver Operating Curve (PR-ROC) to demonstrate difference in the area under the curve (AUC) for the models, RFC and CNN

The features were ranked based on their ability to distinguish between co-occurring and non-co-occurring pairs, with the most important features arranged in decreasing order of their f-scores. Highly ranked features included overrepresentation, PPI score, motif representation features, and DNA shape scores. Among the DNA shape scores, the OC2, EP, and Opening scores were found to be particularly significant based on their high f-score values. (Figure 2).

### 3.2 Both CNN and RFC models perform well and provide their own advantages

The CNN model achieved an accuracy of 94%, while the RFC model achieved an accuracy of 88% in classification. When the *dng* and *dbbs* features were included in the training, the model achieved a very high accuracy of 99% in determining whether binding pairs were co-occurring or not. However, since these features were used to generate the training dataset, they were removed from the final model to avoid overfitting. Instead, other features were used to train the RFC model.

Both the models were evaluated using a precision-recall ROC curve, with the CNN model performing slightly better than the RFC model with an AUC value of 0.94 and 0.88, respectively. The F1 scores were also calculated, with the CNN model achieving 0.938 and the RFC model achieving 0.872. Both models were deemed to have good performance and were used for further predictions on the *discovery set*, with the results available on our webserver.

To predict CRMs, we used the pairs predicted by the neural network as input to a clustering algorithm. The clusters with higher scores are considered to have a higher likelihood of co-occurring and expressing biological significance. Overall, a total of 200k CRMs were generated for 50762 genes, with an average of 10 clusters per gene. Each CRM is typically 300 bp in length and contains binding sites for at least 4 unique TFs, which can contribute up to 10 or more binding sites. The combined cluster score (CRM score) was calculated by averaging the co-occurring scores in a cluster. We output these potential CRMs in BED format for easy visualisation with tools such as IGV browser (Figure S5:d). We provide genome-scale annotation of human CRMs via a web-based database that can be queried with gene or motif names. An automated pipeline for predicting CRMs is available both via a web -interface and as a standalone command-line based version (Figure S6).

## 4 Benchmarking and cardiac CRM identification

### 4.1 We recovered known CRMs identified by previous tools and predicted novel instances

First, we compared our model with MCOT and ChIP-Atlas. We selected motif pairs detected by MCOT and scanned them in our CRMs. We found that the motifs (JUN. USF1), (RXRA, USF1), (AR, FOXA1), and (STAT6, CEBPA) were part of co-localised motif pairs, and were previously reported by [17]. However, Unibind data did not include motif data for IKZF1, and we found a smaller number of binding sites for STAT6. As a result, we were unable to find CRMs that contained these motif pairs. For the remaining three pairs, we found them in our annotated CRMs and obtained additional motifs along with the matched motif pairs (Figure S5: a). In most cases, these pairs co-occurred with other motifs, and we calculated the average CRM size. The counts and CRM sizes are reported in the (Figure S5:a). Similar to our report, ChIP-Atlas also identified the first three pairs, but did not include the interaction between the pairs STAT6-CEBPA and RELAIKZF1. This similarity in the reports may be due to the use of data from the same experimental ChIP-seq data.

Furthermore, we assessed our model’s performance by benchmarking it against a recent CRM identification algorithm known as dePCRM2 [24]. To do this, we compared our CRM predictions with those generated by the dePCRM2 approach using the BEDtools intersection utility. We discovered that all of our CRM predictions exhibited a 100% overlap with the CRMs predicted by dePCRM2. This provided us with a reassuring confirmation that our data-driven methodology effectively identified high-confidence CRMs that had previously been identified by dePCRM2. The latter method accomplishes this by incorporating ChIPseq peaks associated with TF binding, enhancer data, nucleotide variation, and gene expression.

### 4.2 We predict CRMs associated with heart development and diseases

We applied our algorithm using a dataset of known cardiac TFs [39] to predict CRMs. This prediction and analysis are crucial in heart disease research, as subtle alterations in the motif network can influence disease development. CRMs offer insights into the regulatory mechanisms of heart diseases, aiding the development of diagnostic and therapeutic strategies.

Our pipeline predicts CRMs that can be filtered based on type of motifs, genes, or different statistics related to them (Figure S5). Following our definition for CRM we flitered this data to report 1784 Cardiac CRMs with a cut-off score of 0.9 and containing at least four cardiac TFs. A full list of genes and their annotation is provided in the Supplementary Table S1. The TF clusters with critical role in heart function were observed in the data (Figure S5b-d). We narrowed down our results to focus on cardiac TFs associated with genes implicated in a cardiac conditions such as HTAD, SCAD, AVB, AVN, hcCHD, and cardiac dysfunctions (Table 2). Our analysis identified both known and novel TFs. For instance, F11R, a high-priority genetic candidate for SCAD reported in our previous work [41], has a known cardiac TF called PPARG. Our approach enabled us to predict new TFs such as ARID3A and RXRB as potential regulators of SCAD (Table 2). However, the cluster size for this gene falls below our minimum cluster cut-off of four, and prediction of complete clusters may require further analysis of heart specific datasets. Additional exploration revealed that the average cluster size of cardiac CRMs is between 10 and 11 binding sides. The genes with large cluster size include ZCWPW2, ADAL, CDK13, APAF1, and RP11. Moreover, we observed that the REST:JUN, JUN:TP53, and RARA:JUN TF pairs were dominant pairs of cardiac CRMs (Figure 4). Mutations in the JUN and TP53 cardiac motifs have been linked to cardiac failure [45] [46].

**Table 2:**
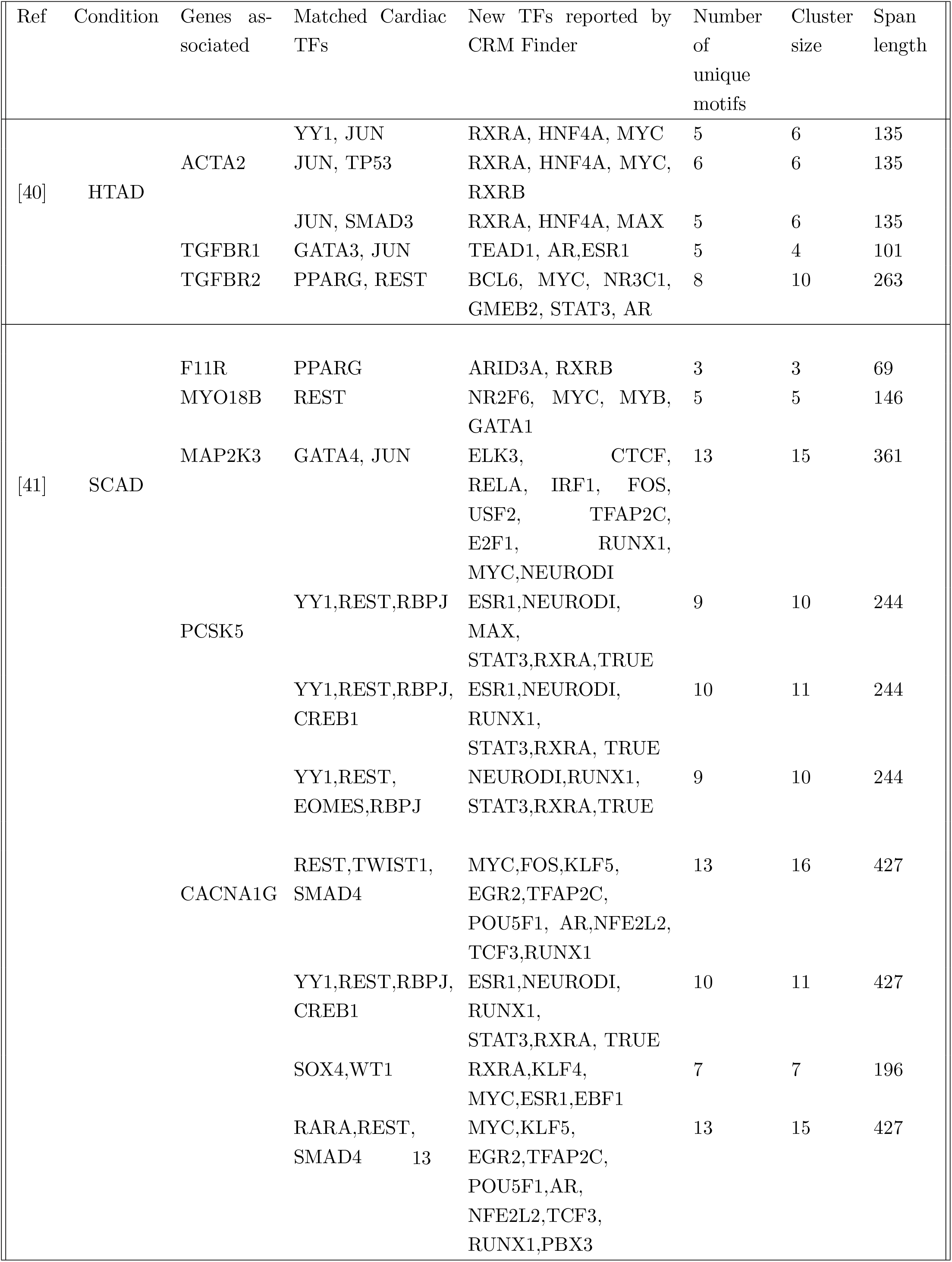
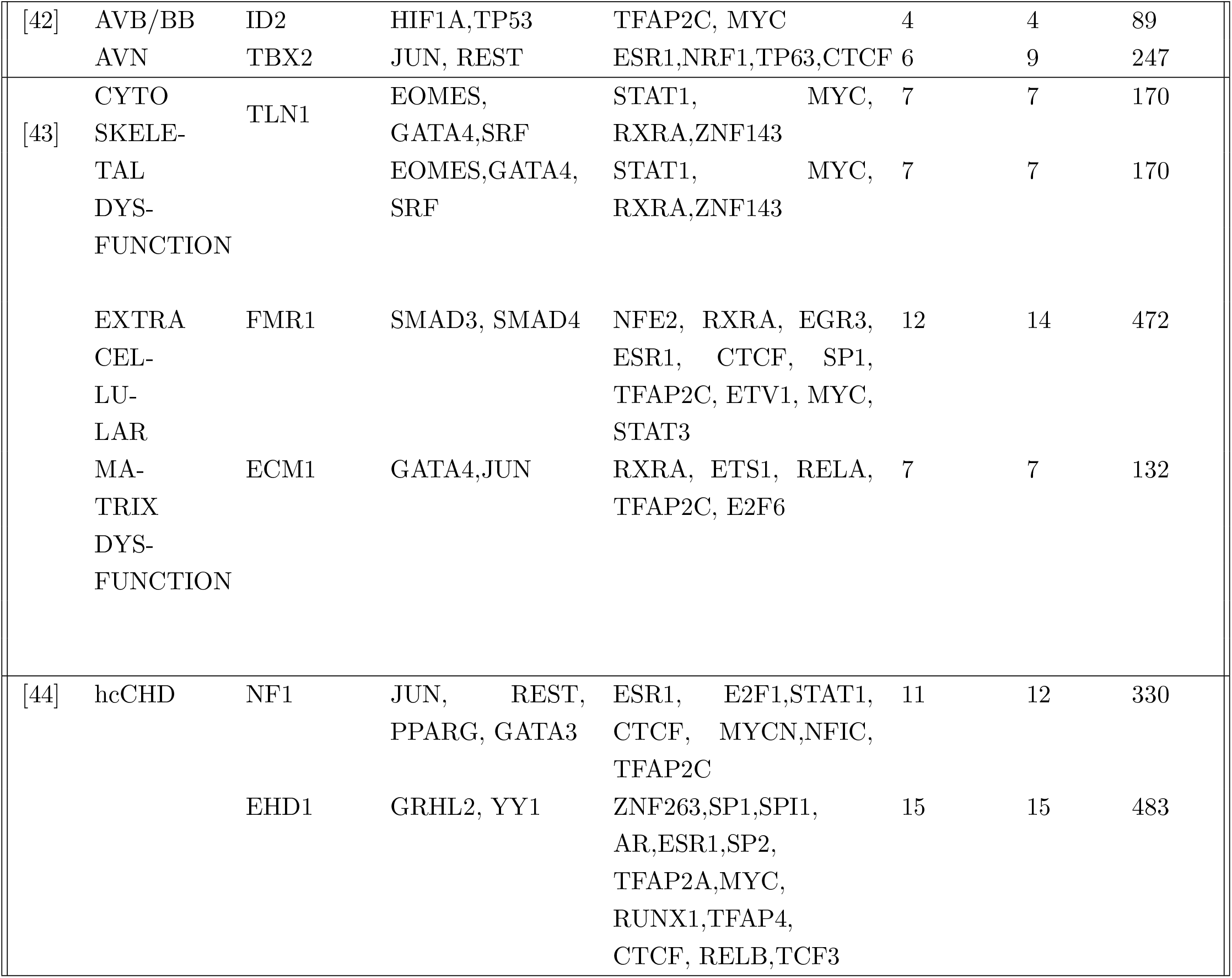
The outputs generated by CRM Finder are matched against existing relevant published work to discover new TFs associated with cardiac conditions.

**Figure 4:**
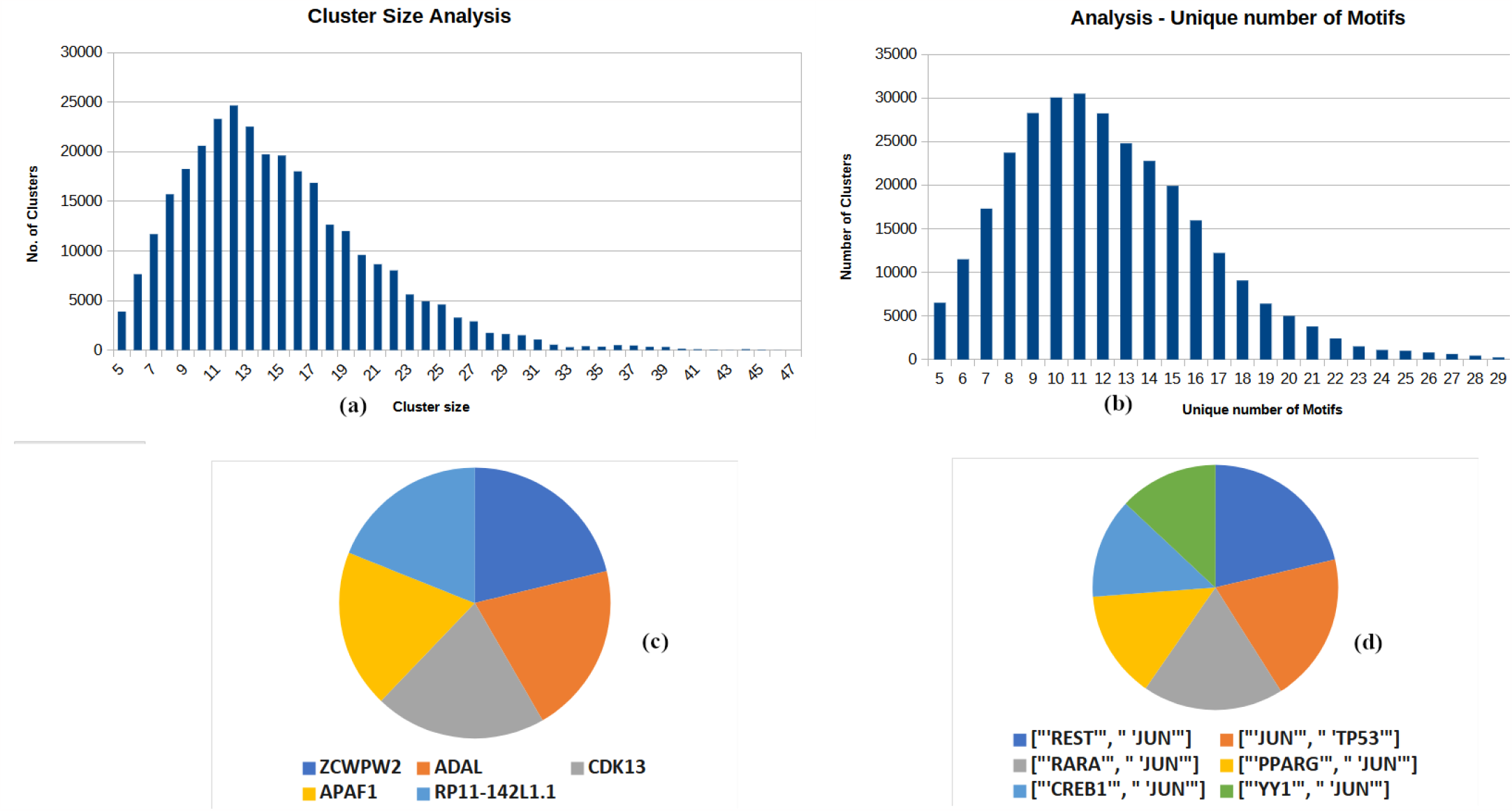
The Cardiac CRMs generated were analysed to get a comprehensive view on the result. The selected CRMs of length greater than four were used for further analysis. (a) This shows how many clusters are reported under each cluster size and we find most of the clusters have size 10 or 11.(b) The most frequent cluster size after considering only unique motifs was found to be between 11 and 15 (c) The genes which have most number of TFs are depicted here (d) Frequent co-occuring cardiac TFs are plotted here

### 4.3 Cardiac gene regulatory networks

The role of the NKX family of TFs in cardiac development [39] was investigated by examining a subset of cardiac CRMs that contained at least one NKX TF binding site. Network analysis (Figure 5a) and K-mean clustering of genes containing these TFs revealed three interaction clusters (reported as red, blue, and green in the Table S2), and the top cluster (red) from here displayed significant enrichment for genes associated with expression, diseases, and biological pathways that are specific to cardiac tissues (Figure 5b). These findings suggest that the NKX TF family may play an important role in regulating the cardiac gene regulatory network, and highlight potential targets for further investigation in cardiac development and disease.

**Figure 5:**
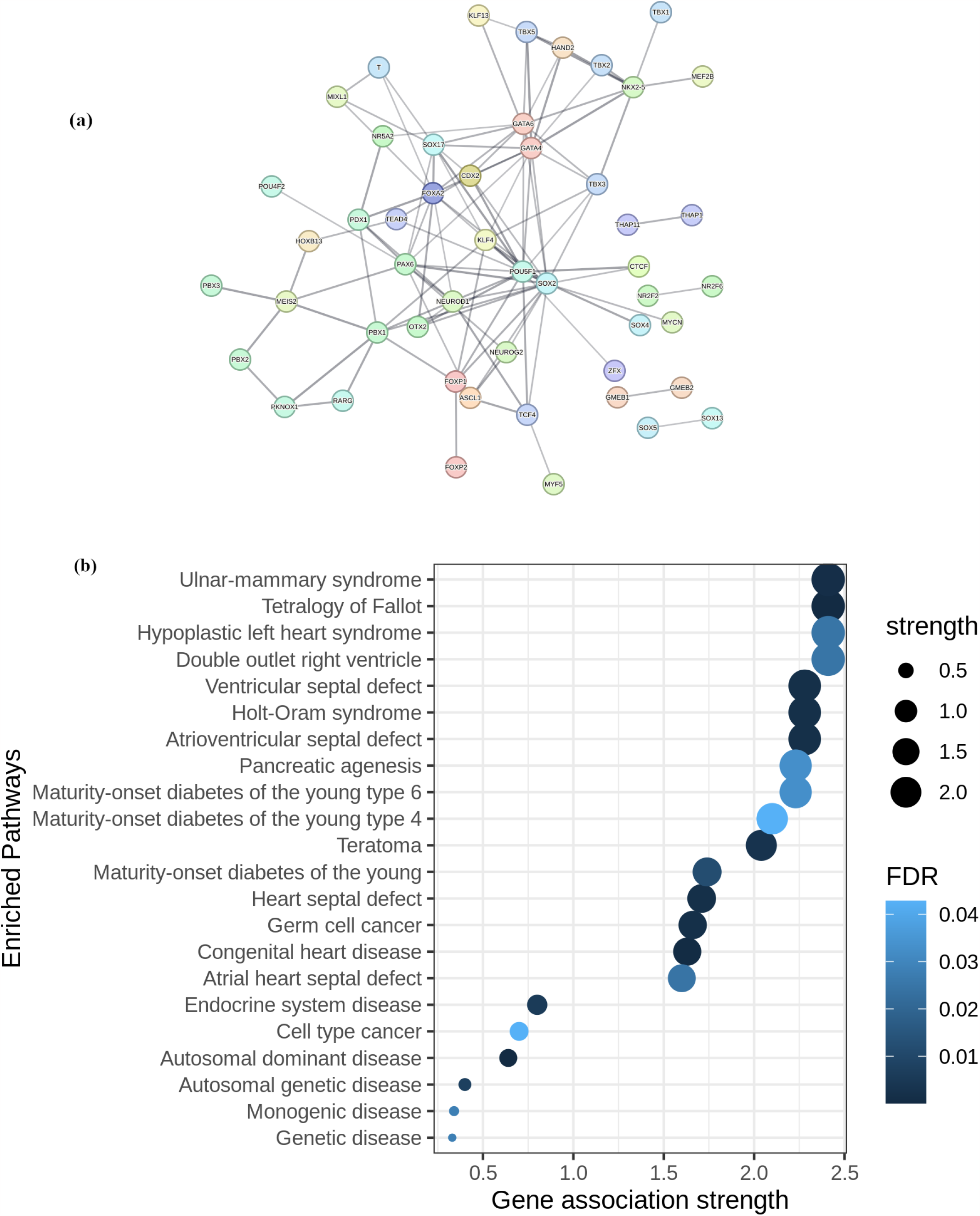
Cardiac CRM analysis using the STRING app [26]: (a) Interaction network of CRMs containing one of NKX family TFs. (b) The enrichment analysis of the NKX-TF red cluster shows strong association with heart disorders.

Furthermore, to determine the efficacy of our pipeline, we investigated whether it contained the genes networks as reported by Kiran Musunuru *et al* [40][47]. Our analysis revealed that we were able to identify a subset of genes with strong evidence for HTAD (heritable thoracic aortic aneurysm or dissection) in our pipeline results. We then extracted network of various TFs that were identified for all of these genes, as shown in the Figure 6.

**Figure 6:**
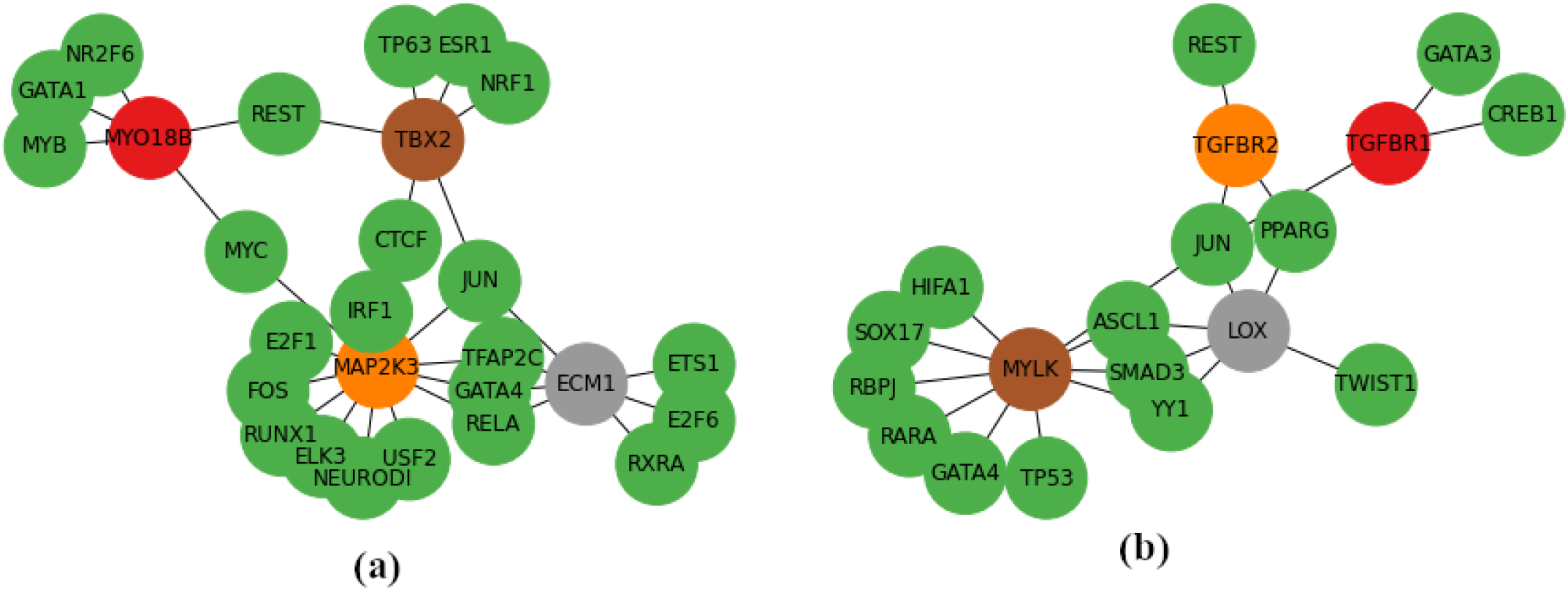
GRNs reported by our pipeline: (a) Depicts a regulatory network where green nodes represent TFs and other colored nodes represent genes. Here, genes MYO18B, MAP2K3, ECM1,TBX2 presented CRMs with common TFs. MYO18B, MAP2K3 are associated with SCAD whereas, genes ECM1, TBX2 are responsible for arterial matrix dysfunction and cardiac networks development. [42] [43] (b) Here, genes TGFBR2, TGFBR1, LOX, MYLK have common TFs and were previously reported to be associated with HTAD condition [40]

## 5 Data and Software Availability

The novel CRMs that our pipeline generates can be accessed through our web-hosted database https://bioinformaticslab.erc.monash.edu/CRMdatabase; retrieved on 10 October 2023; see Figure S6). Our pipeline implements the latest software and distributed environment strategies, allowing for the efficient processing of large genomic datasets. The source code for our pipeline is readily available for public use (https://gitlab.com/tyagilab/crc_finder); retrieved on 10 October 2023.

## 6 Conclusion

Our study offers a comprehensive survey of current CRM-finding approaches and addresses significant gaps in the field. We introduce a novel pipeline for predicting putative CRMs from identified DNA motifs, which can facilitate the study of gene regulation and motif interactions. Our pipeline incorporates both RFC and CNN machine learning models for predicting TF co-occurrence and clustering to identify new CRMs. We also provide a complete annotation of 223 human motifs (excluding the sites used in model training) from the UniBind database, which is accessible through our user-friendly web interface. Our pipeline is highly versatile and can be run via our web server or a command-line interface.

This work identified several important factors in predicting TFBS cooperativity, including TFBS overrepresentation, PPI score, motif representation features, and DNA shape scores. In particular, the OC2, EP, and Opening scores (shape features) were found to be highly significant. Using this pipeline, we were able to discover novel cardiac CRMs and GRNs that are associated with cardiac development and disease pathways. The versatility of this pipeline allows it to be applied to a wide range of biological systems and datasets beyond the cardiac development and disease pathways studied in this work. By leveraging the factors identified for predicting TFBS cooperativity, this approach has the potential to uncover novel regulatory pathways involved in various disease and developmental processes. Our approach is species-agnostic and highly flexible meaning it could be extended to other large-scale datasets from similar assays, such as ATAC-seq, ChIP-HiC, or DNAse-seq.

## 7 Acknowledgements

ST acknowledges support from Australian Academy of Science, Australian Women Research Success Grant at Monash University. We acknowledge the helpful discussions and compute resources from the Monash eResearch Platform and the MASSIVE HPC facility (www.massive.org.au). We thank Yashpal Ramakrishanaiah and Tyrone Chen for their feedback and help in packagaing the software and setting up the webserver.

## 8 Supplementary Data

### 8.1 Co-binding distance

**Figure S1:**
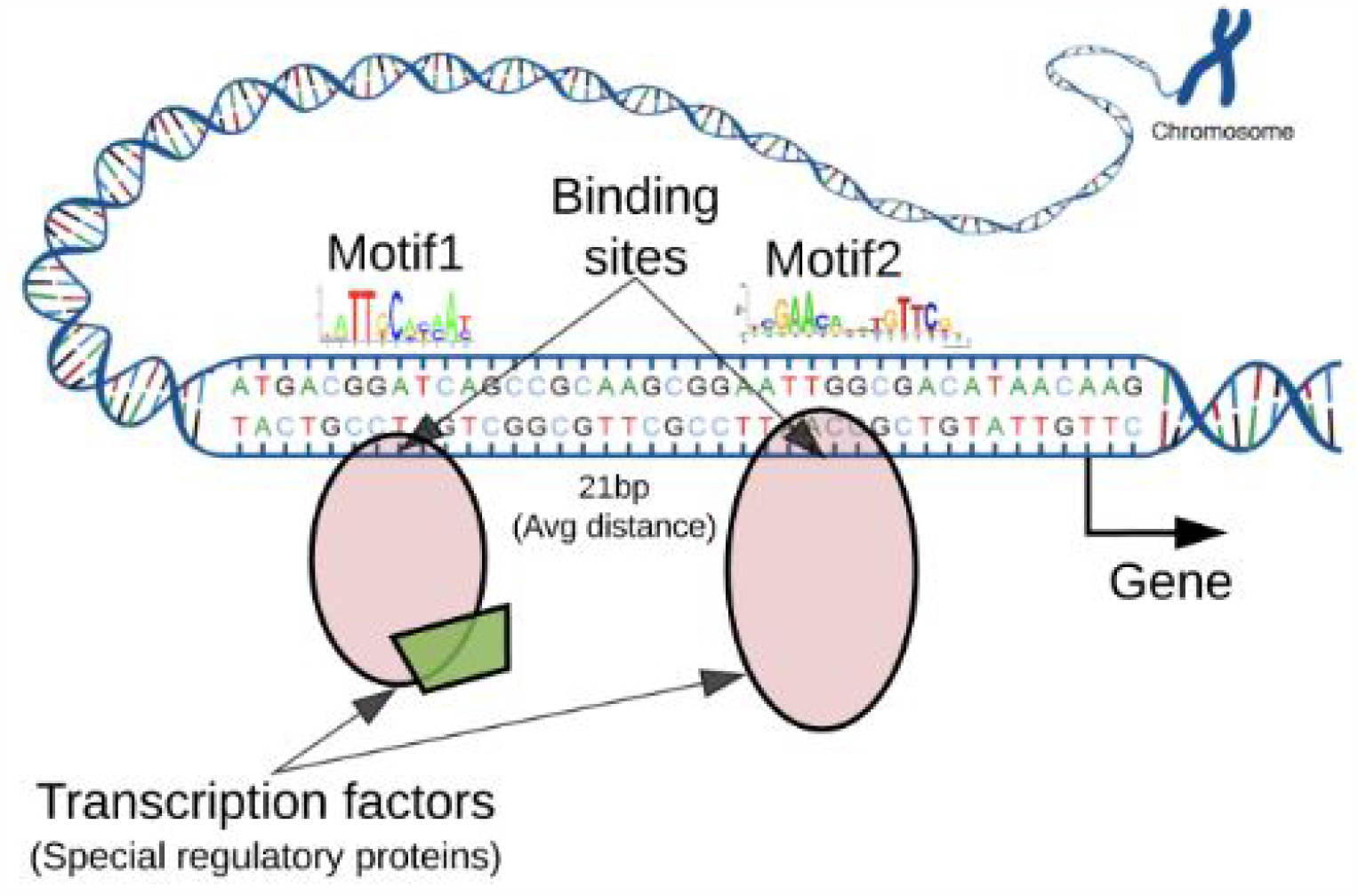
Generally millions of binding sites for a TF are found but only a few thousands are bound. GATA1 and TAL1 are a known genome-wide co-occurring motif pair. This pair is known to cooperate to regulate erythroid development across thousands of genomic locations on the DNA

**Figure S2:**
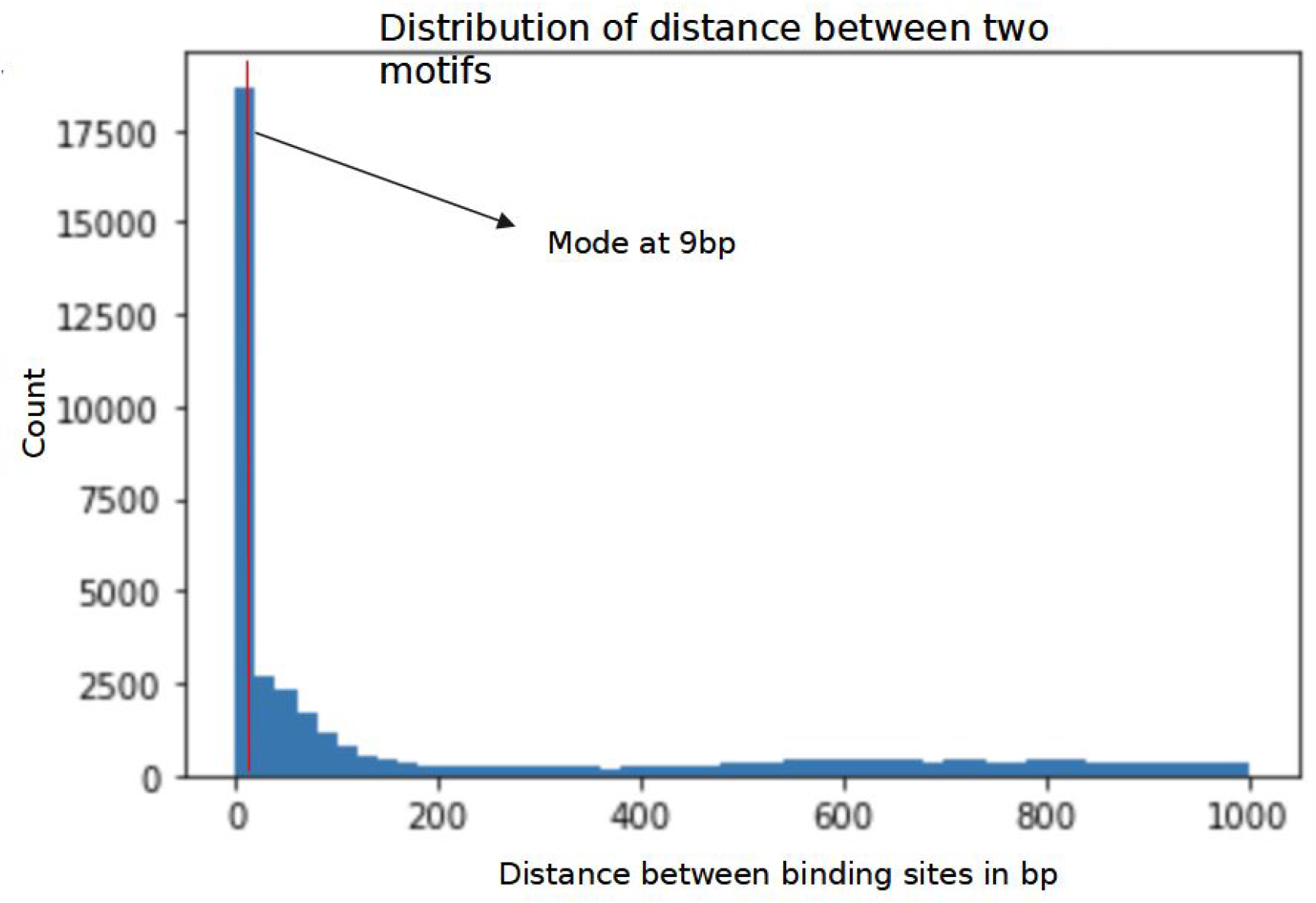
Distance between binding sites (*dbbs*) in the promoter region shows a mode at 9 bp. Previously, an average of 21 bp distance was observed [29]. We relaxed this distance to be 30 bp in our methods to allow for any unknown variations.

**Figure S3:**
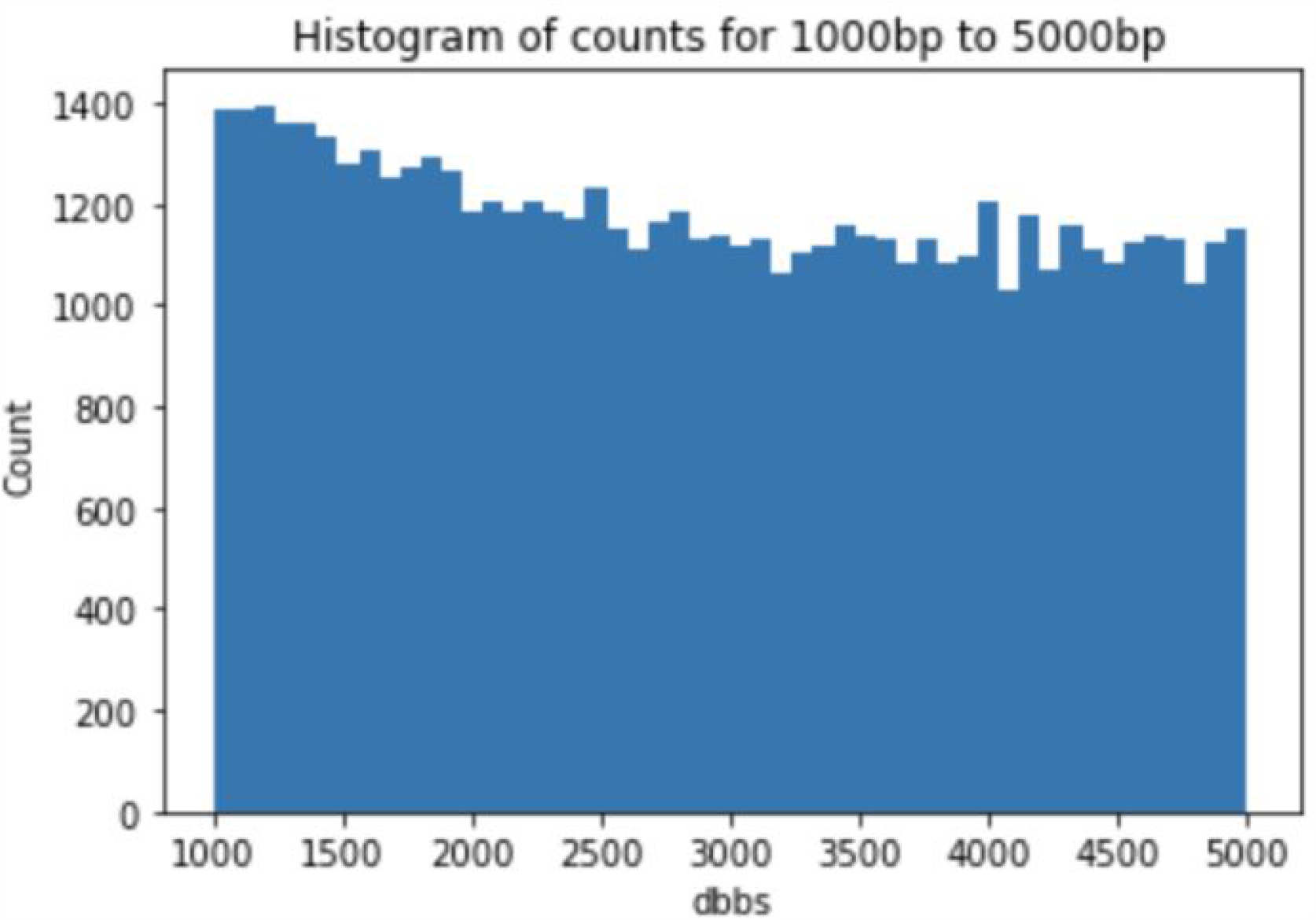
Motif Pairs of distances greater than 1500 bp shows a uniform distribution(Figure S3).

### 8.2 Feature Ranking

To calculate and quantify the feature importance, we performed univariate feature importance tests. These tests involved the Chi-square test and one-way ANOVA test (f-score). These univariate methods do not consider any classification algorithm for feature importance. Chisquare is a goodness of fit test which is performed to check if the sample belongs to a population which returns a chi-square score. The f-score indicates the difference between the distributions of values between the classes. The chi-square (*χ*^2^) and the f-scores (F) were calculated for each of the features against the target variable with the following formulae.

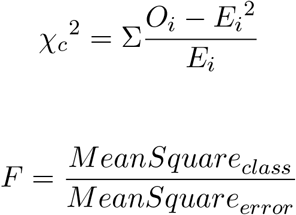

Recursive Feature Elimination with Cross-Validation was performed to select the most important features for data classification. This technique involves the iterative elimination of the least important feature and the importance scores are recalculated. As observed in the (Figure S4), 9 features were sufficient for optimal model training.

**Figure S4:**
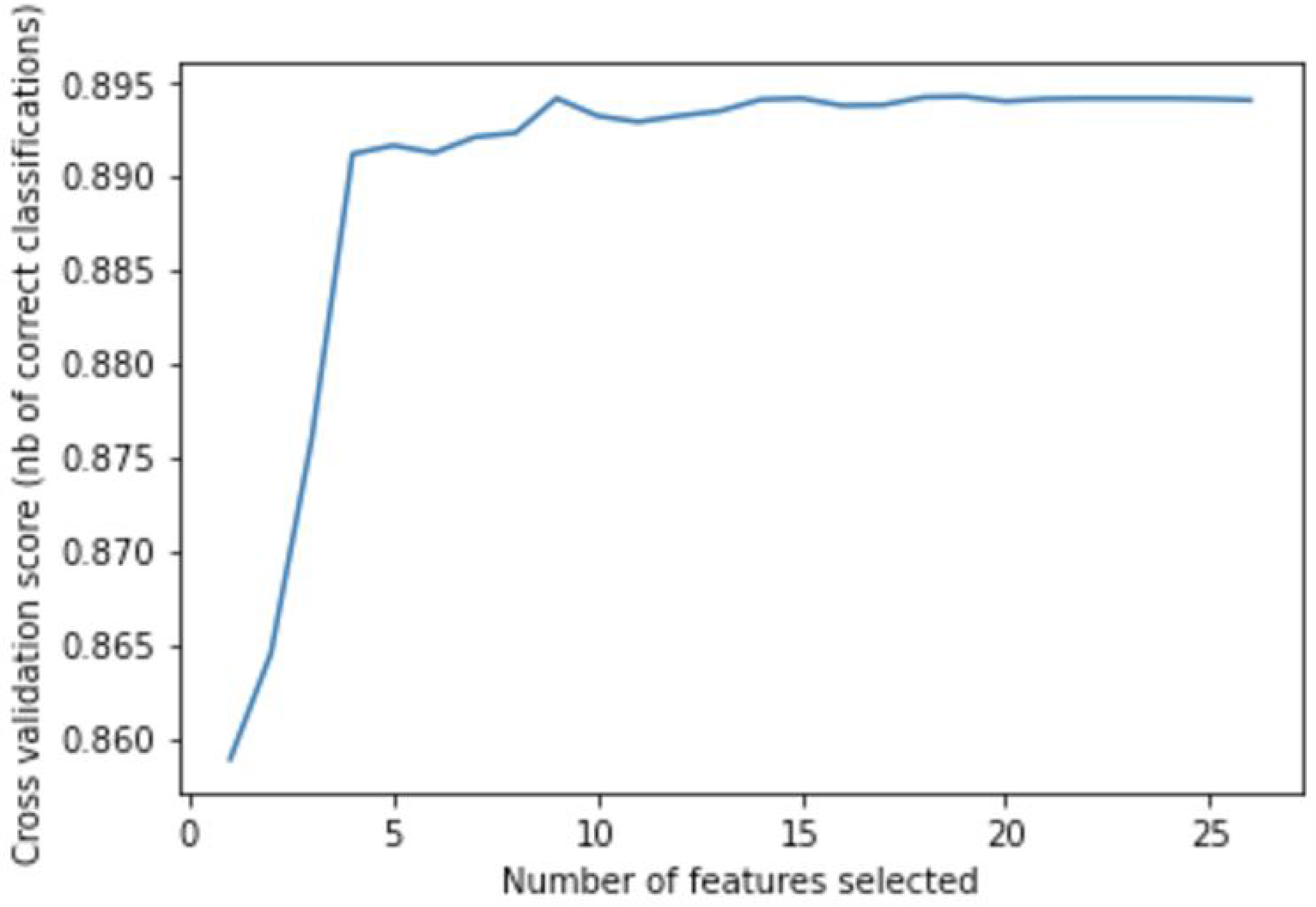
Recursive Feature Elimination with Cross-Validation (RFECV)

### 8.3 Predictions of Cardiac CRMs and benchmarking

We filtered cardiac CRMs of atleast 1 unique cardiac TFs with clustering score greater than 0.9 which resulted in approximately 150K CRMs (Figure S5). The gene list generated from the predicted cardiac CRMs was analysed. Their gene type, GO term name and annotation can be found in the Table S1.

[Table S1: filename=Cardiac CRM genes and types.txt]

Table S2: filename=NKX-cluster-analysis.xlsx

### 8.4 CRM Website

Using the web application, users can extract CRMs for specific genes. We also provide options to set criteria like extract using specific motifs or known PPI score of TFs and cluster score cut off. The results can be downloaded as CSV

**Figure S5:**
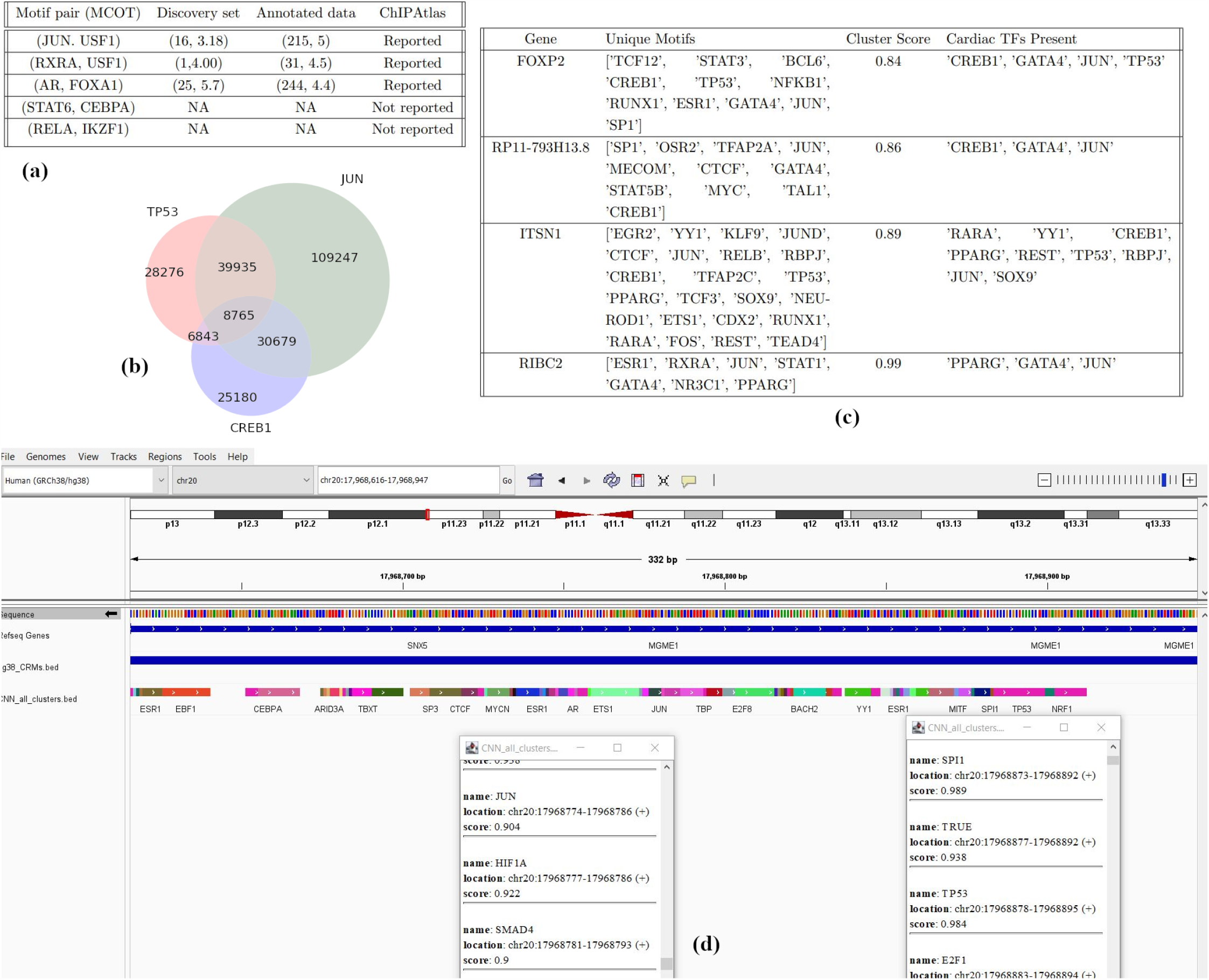
(a) We evaluated the accuracy of our predictions by comparing them to data from MCOT and ChIP Atlas. The validation results are presented. (b) We examined the co-occurrence of TP53, a TF with important role in the functioning of heart. The results are depicted in a Venn diagram, where CREB1 and JUN were the most common TFs forming clusters with TP53. (c) A sample output shows the genes with at least one cardiac TFs listed in their clusters. (d) IGV window showing cardiac TFs TP53, JUN, HIFA1, SMAD4, SPI1, E2F1 with a CRM score of 0.92.

**Figure S6:**
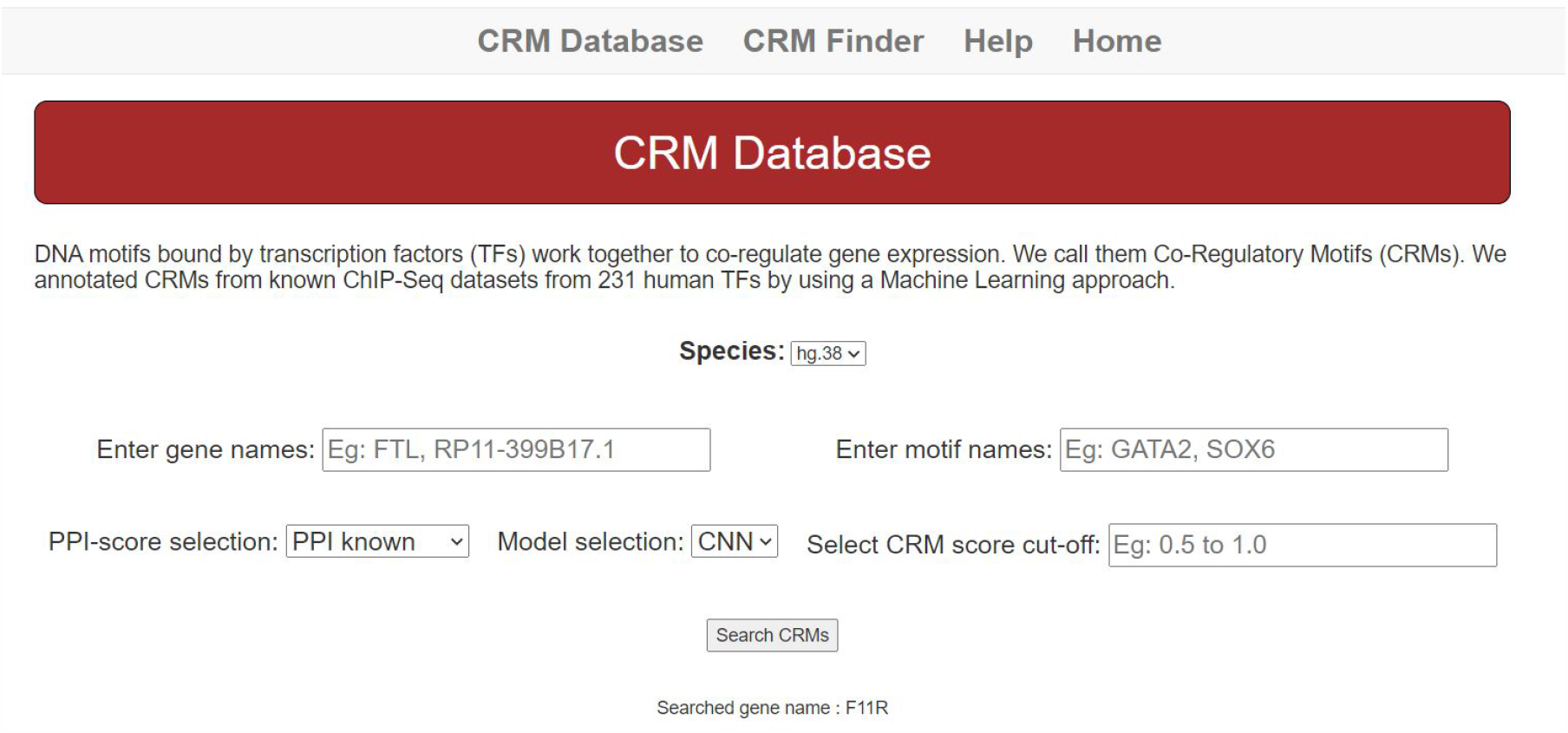
CRM finder and an annotated database of human motifs is accessible via a Webserver

